# High-throughput CRISPR live-cell imaging of low-frequency chromosomal events quantifies the latent efficiency of chromosome engineering

**DOI:** 10.64898/2026.05.10.724155

**Authors:** Xixun Hu, Yuichiro Iwamoto, Kyotaro Yamazaki, Nanami Kishima, Natsuko Otaki, Hitomaru Miyamoto, Yuichiro Miyaoka, Yasuhiro Kazuki, Sadao Ota

**Affiliations:** Research Center for Advanced Science and Technology, The University of Tokyo, 4-6-1 Komaba, Meguro-ku, Tokyo, 153–8904 Japan; Exploratory Research Center on Life and Living Systems (ExCELLS), National Institutes of Natural Sciences, 5-1 Higashiyama, Myodaiji-cho, Okazaki, Aichi 444-8787 Japan; National Institute for Physiological Sciences, National Institutes of Natural Sciences, 5-1 Higashiyama, Myodaiji-cho, Okazaki, Aichi 444-8787 Japan; Department of Chromosome Biomedical Engineering, Integrated Medical Sciences, Graduate School of Medical Sciences, Tottori University, 86 Nishi-cho, Yonago, Tottori 683-8503, Japan; Artificial Intelligence Medicine, Graduate School of Medicine, Chiba University, 1-8-1 Inohana, Chuo-ku, Chiba City, Chiba 260-8670, Japan; Institute for Advanced Academic Research (IAAR), Chiba University, 1-33 Yayoi-Cho, Inage-ku, Chiba City, Chiba 263-8522, Japan; Regenerative Medicine Project, Tokyo Metropolitan Institute of Medical Science, Setagaya, Tokyo, 156-8506, Japan; Graduate School of Medical and Dental Sciences, Institute of Science Tokyo, Bunkyo, Tokyo, 113-8510, Japan; Graduate School of Humanities and Sciences, Ochanomizu University, Bunkyo, 112- 8610, Tokyo, Japan; Chromosome Engineering Research Center, Tottori University, 86 Nishi-cho, Yonago, Tottori 683-8503, Japan

## Abstract

Quantifying low-frequency chromosomal alterations in living cell populations at early stages is essential in many fields including cancer studies and chromosome engineering, yet selection-based readouts impose delays and can lose fragile positives before readout, biasing frequency estimates; CRISPR imaging rarely reports detection limits at 10^−4. Here, we developed High-throughput CRISPR Imaging (Hi-CRI), integrating engineered dCas9–sgRNA ribonucleoprotein (RNP) labeling, suppression of nonspecific aggregates via metabolic modulation and protease treatment, high-speed volumetric imaging by oblique plane microscopy, GPU-accelerated image analysis, and an explicit error-controlled detection-limit framework. Using per-cell signal-to-noise ratio calling, Hi-CRI achieves a 0.01% detection limit for target-positive cell fractions. In microcell-mediated chromosome transfer of a mouse artificial chromosome (MAC) into HT1080 recipients, Hi-CRI measured 0.03% MAC-positive cells among 184,235 recipients at day 1 post-fusion, versus 0.0007% by antibiotic-selection-based clonogenic assay at day 8 post-fusion, consistent with substantial loss before readout (pre-readout attrition). Hi-CRI enables viability-preserving, selection-independent quantification of low-frequency chromosomal states.

**Teaser:** Rare chromosome events can be counted in living cells by high-throughput CRISPR imaging before selection hides them.

## Introduction

Chromosomal alterations—ranging from aneuploidy and polyploidy to targeted rearrangements—are key sources of cellular heterogeneity that shape the evolutionary trajectories of cancer *(1–5*) and underpin diverse strategies in chromosome engineering *(6–9)*. Understanding the onset and dynamics of these alterations requires the ability to detect and track rare cells carrying specific chromosomal changes within large populations. In cancer, for instance, the emergence of rare whole-genome-doubled subclones can precede tumor progression and drug resistance *(5)*. In chromosome engineering contexts such as microcell-mediated chromosome transfer (MMCT), successful delivery of exogenous chromosomes likewise occurs at low frequency *(7)*, leaving modified cells obscured within a large excess of unmodified cells.

However, a critical methodological gap limits accurate quantification of such rare chromosomal events. Conventional detection typically relies on retrospective selection, such as antibiotic resistance or prolonged clonal expansion, which imposes severe survival pressure and a substantial temporal delay. As a result, fragile, newly engineered subpopulations are preferentially lost due to antibiotic toxicity, metabolic stress, or transient chromosomal instability before colony formation *(7)*. We refer to this cumulative loss before measurement as pre-readout attrition, which can obscure the frequency of chromosomal events in the starting population.

Imaging in living cells offers a non-destructive alternative, but existing CRISPR-based labeling approaches *(10–13)* lack the combination of sensitivity and throughput required to detect events at frequencies of 10^−4^ or lower. The signal-to-noise ratio (SNR) is often limited by nonspecific aggregation of CRISPR components *(14)*, and conventional three-dimensional microscopies do not readily scale to populations of 10^4^-10^5^ cells (with sufficient statistical confidence). Moreover, CRISPR imaging has not widely adopted a formal detection-limit framework with explicit error control, aligned with established definitions, making frequency claims near 10^−4^ difficult to substantiate and compare across studies.

To address these limitations, we developed a high-throughput CRISPR Imaging (Hi-CRI) platform that integrates oblique plane microscopy *(15–17)* and GPU-accelerated volumetric analysis, and an optimized CRISPR-based labeling workflow. Hi-CRI combines engineered dCas9 ribonucleoproteins (RNPs) with metabolic modulation to suppress nonspecific nuclear RNP aggregations, achieving a limit of detection (LoD) of 0.01% under definitions from the International Union of Pure and Applied Chemistry (IUPAC). When applied to MMCT, Hi-CRI measured a 0.03% chromosome transfer efficiency, which is substantially higher than the estimates obtained by conventional antibiotic selection. This discrepancy is consistent with the substantial pre-readout attrition inherent to selection-based assays, indicating that Hi-CRI captures a larger fraction of chromosome transfer events that would otherwise be lost before readout. Together, these results establish Hi-CRI as a broadly applicable framework for direct, viability-preserving quantification of low-frequency genomic events.

## Results

### An integrated workflow for detecting low-frequency target-chromosome-positive cells in living populations

Figure 1 depicts the overview of Hi-CRI workflow. To label target chromosomes in live cells, we first delivered engineered fluorescent-protein-tagged dCas9 RNP complexes into cells by electroporation (Fig. 1A). dCas9 RNPs programmed with single-guide RNAs (sgRNA) accumulate specifically at the target locus and form punctate nuclear foci, hereafter referred to as CRISPR spots. To minimize spurious foci and improve detection at low frequencies, as detailed later, we engineered the RNP and refined the labeling workflow to suppress nonspecific nuclear RNP aggregation.

**Fig. 1.**
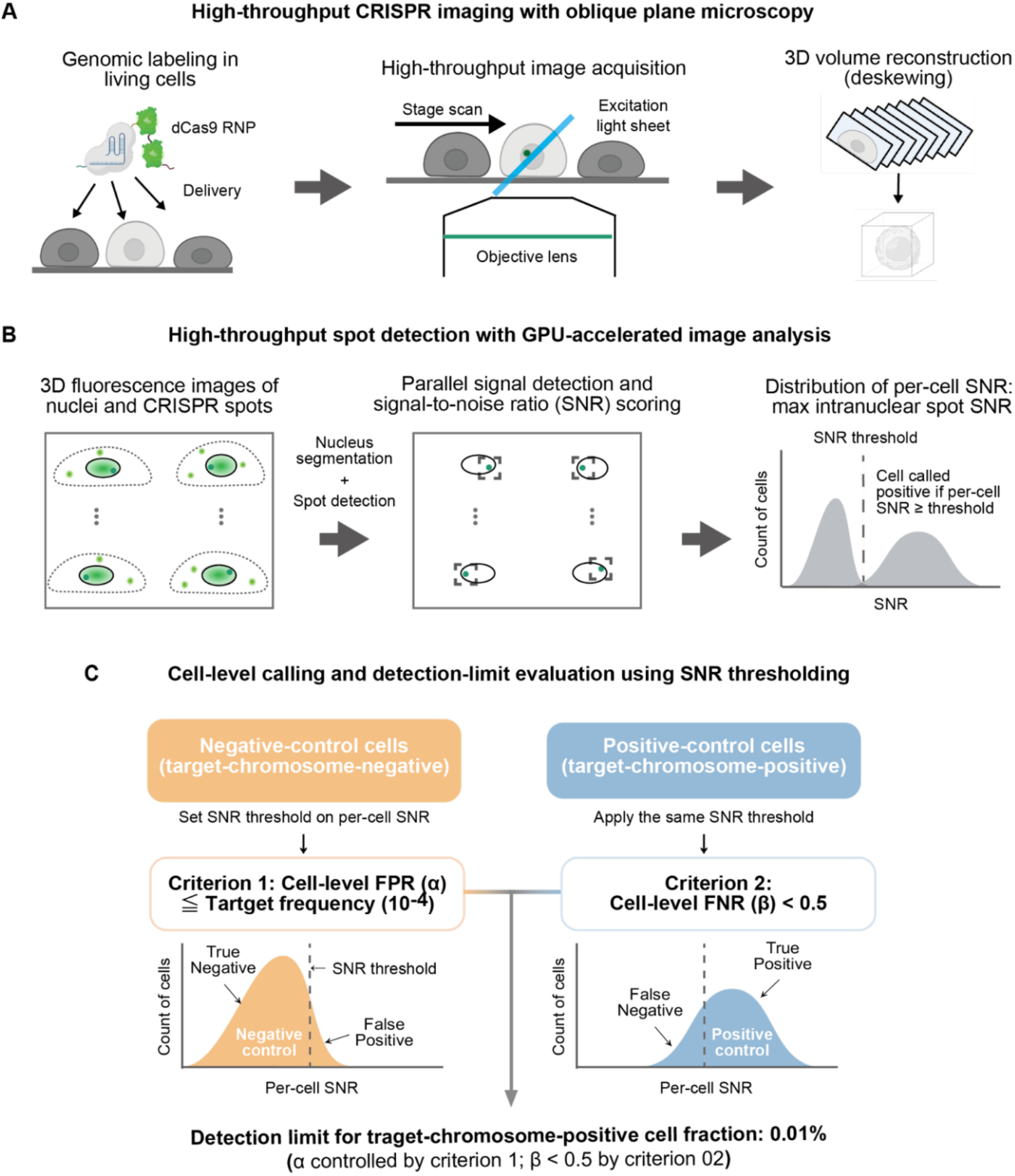
Overview of Hi-CRI, an end-to-end workflow for detecting low-frequency target-chromosome positive cells in living populations. (A) High-throughput CRISPR imaging with oblique plane microscopy. Target chromosomes are labeled in living cells by electroporation of dCas9–GFP ribonucleoprotein (RNP) complexes, followed by high-throughput volumetric imaging using oblique plane microscopy and GPU-based 3D volume reconstruction. (B) High-throughput spot detection with GPU-accelerated image analysis. Three-dimensional nuclear segmentation is performed on reconstructed volumes, followed by parallel CRISPR spot detection, and per-spot signal-to-noise ratio (SNR) scoring. Per-cell SNR is defined as the maximum intranuclear spot SNR within each nucleus, yielding a per-cell SNR distribution. (C) Cell-level signal calling and detection-limit evaluation using SNR thresholding. Cells are called positive if at least one intranuclear spot exceeds the SNR threshold (equivalently, if the per-cell SNR exceeds the threshold). The threshold is set on negative controls to control the cell-level false positive rate (FPR, α) relative to a target frequency (criterion 1) and validated on positive controls by a predefined false negative rate (FNR, β) criterion (criterion 2). Cell-level FPR and FNR are defined as the fractions of target-negative and target-positive control cells, respectively, that are misclassified under the SNR threshold.

To capture CRISPR spots at a population scale, we employed high-throughput oblique-plane microscopy (OPM), a single-objective light-sheet technique, avoiding the physical interference posed by high-NA objectives and enabling high-throughput volumetric imaging (Fig. 1A). Because OPM data are acquired in a sheared coordinate system, raw stacks were deskewed and resampled using a GPU-accelerated affine transformation at 0.2 gigavoxels per second (Fig. 1A).

We next constructed a GPU-accelerated analysis pipeline for per-cell calling. First, nuclei were segmented using Voronoi–Otsu labeling to restrict analysis to intranuclear regions, and spot candidates were detected by centroid-localization–based peak finding (*18, 19*). For each candidate spot, the signal-to-noise ratio (SNR) was computed as the spot intensity divided by the standard deviation of intensities in a local intranuclear background region; the maximum spot SNR within each nucleus was then used as the per-cell SNR for downstream thresholding, yielding per-cell SNR distributions (Fig. 1B). Parallelized segmentation and per-nucleus spot detection achieved an analysis throughput of 29.31 cells per second.

To support quantitative claims at frequencies of ≤ 0.01%, we evaluated a detection limit for the target-(chromosome)-positive cell fraction using an IUPAC-aligned hypothesis-testing framework with explicit control of false positives (α) and false negatives (β) (*20*) (Fig. 1C). A cell was called positive if its per-cell SNR exceeded a threshold. We first set the SNR threshold using only target-negative control cells such that the cell-level false positive rate (FPR, corresponding to α) did not exceed the target frequency of 0.01% (criterion 1). We then applied the same threshold to target-positive control cells and verified that the cell-level false negative rate (FNR, corresponding to β) satisfied the predefined sensitivity criterion used here (FNR < 0.5, corresponding to >50% detection probability) (criterion 2). Under these criteria and acquisition conditions, Hi-CRI meets the target detection limit of 0.01% for the target-positive cell fraction.

### Engineering a recombinant dCas9 RNP for sensitive and specific chromosome labeling in living cells

To reduce false-positives while maintaining high on-target labeling efficiency required for low-frequency chromosome detection, we used an RNP-based labeling strategy that delivers preassembled dCas9–sgRNA complexes into live cells by electroporation. This format enables titratable dosing and reduces cell-to-cell variability relative to expression-based delivery.

To identify a robust RNP design suitable for sensitive chromosome labeling, we systematically optimized two elements of the fluorescent dCas9 fusion protein: nuclear localization signal (NLS) and the fluorescent protein configurations. Optimization of NLS position and copy number revealed that three tandem 2× nuclear localization signal (NLS) modules positioned at the N terminus, an internal site, and the C terminus produced the strongest nuclear enrichment (Fig. S1). Optimization of the number and configuration of fluorescent proteins showed that an N-terminal sfGFP–mNeonGreen tandem fusion produced the brightest CRISPR spots (Fig. S2). We therefore used dCas9–sfGFP– mNeonGreen with three 2×NLSs (hereafter dCas9–green) as the final design for all subsequent experiments (Fig. 2A; Table S1).

**Fig. 2.**
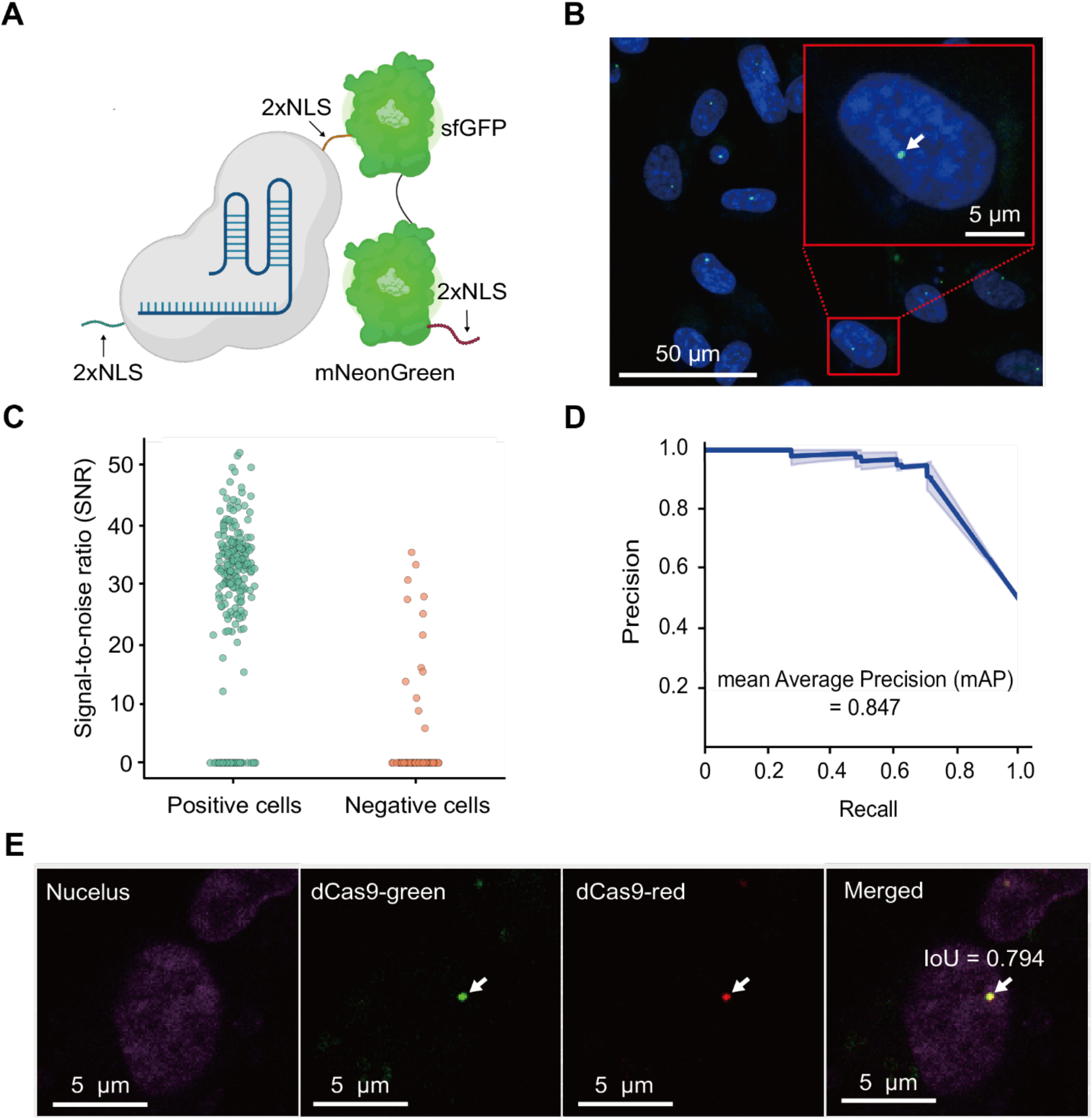
Engineering fluorescent dCas9 ribonucleoproteins for sensitive and specific chromosome labeling in living cells. (A) Schematic of the recombinant dCas9 used for chromosome labeling in live cells. Two green fluorophores (sfGFP and mNeonGreen) are fused at the N terminus, and three tandem 2× nuclear localization signal (NLS) modules are positioned at the N terminus, an internal site, and the C terminus to enhance nuclear import. (B) Representative labeling of a mouse artificial chromosome (MAC) in an HT1080 MAC-positive line imaged by confocal microscopy. Nuclei are stained with Hoechst 33342. The inset highlights a representative CRISPR spot (arrow). Scale bars, 50 µm (overview) and 5 µm (inset). (C) Per-cell signal-to-noise ratio (SNR) distribution for MAC-positive and MAC-negative HT1080 cell lines (n = 262 MAC-positive cells and n = 160 MAC-negative cells across three independent experiments). Per-cell SNR was defined as the maximum intranuclear spot SNR within each nucleus. For cells with no detectable spot, the SNR was set to 0. (D) Precision–recall (PR) performance for MAC detection obtained by sweeping the SNR threshold. The curve shows mean performance across three independent experiments; the shaded band indicates ± standard deviation (s.d.); mAP is mean average precision. (E) Two-color colocalization assay using dCas9–green (sfGFP–mNeonGreen) and dCas9– red (2×mScarlet) programmed with distinct guide sets targeting the MAC (sgMajSat_1 for green and sgMajSat_2 for red). Nuclei are stained using NucSpot Live 650. The inset highlights a representative CRISPR spot of MAC focus (arrow). Colocalization was assessed by intersection-over-union (IoU) between the green and red focus masks, defined as the area of overlap divided by the area of union; spots were classified as colocalized at IoU ≥ 0.5. Scale bars, 5 μm.

To evaluate live-cell chromosome labeling using the recombinant dCas9-green, we pre-assembled dCas9–sgRNA RNPs targeting the mouse artificial chromosome (MAC) or human chromosome 19 and delivered them by electroporation. Discrete nuclear CRISPR spots became detectable approximately 8 h after delivery under our culture conditions, including in HT1080 cells (Fig. S3). We first quantified labeling performance in isogenic HT1080 lines with and without a single-copy MAC. Guides targeting mouse major satellite (MajSat) repeats produced bright punctate signals in the MAC-positive line (Fig. 2B). We quantified per-cell SNR, defined as the maximum intranuclear spot SNR within each nucleus, and observed distinct distribution between MAC-positive and MAC-negative HT1080 controls (Fig. 2C), consistent with on-target accumulation at the labeled locus. Using the MAC-positive and MAC-negative lines as ground-truth controls, we computed precision and recall across SNR thresholds to generate a precision–recall curve, yielding a mean average precision (mAP) of 0.847 (Fig. 2D). To further assess specificity, we performed two-color colocalization using dCas9–green (sfGFP–mNeonGreen) and dCas9–red (2×mScarlet) programmed with distinct guide sets targeting separate MAC loci (Table S2; Data S1). We quantified colocalization using intersection-over-union (IoU), a mask-overlap metric computed between the green and red foci (Fig. 2E), showing that 96.0% ± 0.5% of spots were colocalized at IoU ≥ 0.5 (Fig. 2E; Table S3).

In addition, we qualitatively confirmed that CRISPR spots were detectable with the same workflow across additional cell lines (U2OS, K562, Jurkat, and CHO) and for an additional genomic target (human chromosome 19) (Fig. S3).

Together, these results establish an RNP implementation that supports sensitive and specific chromosome labeling in living cells, providing a foundation for subsequent high-throughput imaging and quantitative signal calling.

### Optimizing the labeling workflow via metabolic modulation to suppress nonspecific aggregates

With an engineered fluorescent dCas9 RNP that supports sensitive and specific chromosome labeling in living cells (Fig. 2), we next asked whether the full assay could meet the detection-limits required to quantify low-frequency events in large populations. Because evaluation at target frequencies near 10^−4 requires readout across large cohorts, we benchmarked detection limits using the high-throughput volumetric imaging and GPU-based analysis pipeline described above (Fig. 1). Under the initial labeling workflow, negative-control cells exhibited a distinct high-SNR tail consistent with nonspecific puncta, and detection-limit analysis yielded an LoD of approximately 0.1% under the defined acquisition and calling procedure (Fig. 3D). This result indicated that workflow-level suppression of nonspecific signals, rather than RNP design, had become the limiting factor for reaching the 10^−4 regime.

**Fig. 3.**
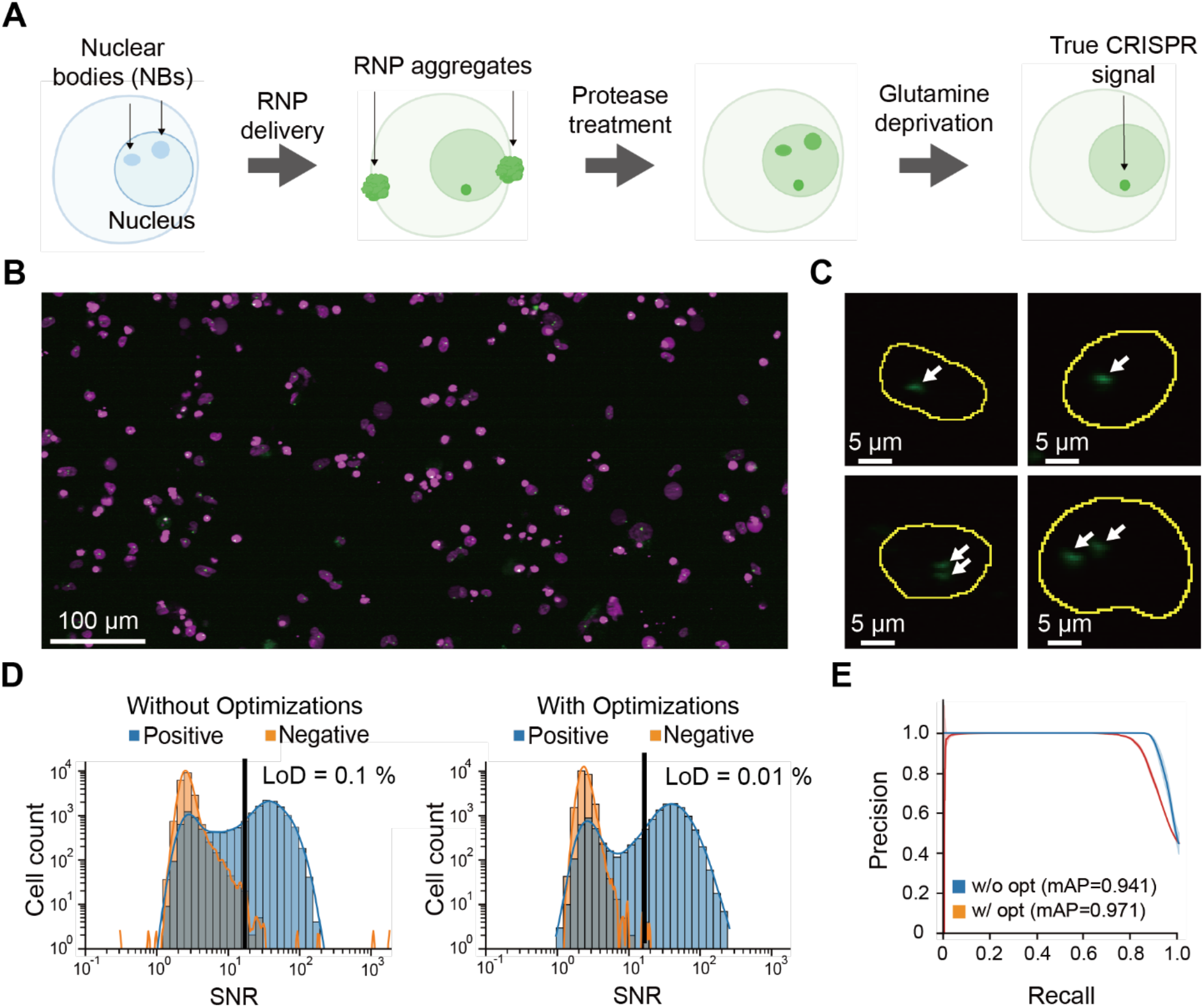
Workflow optimization suppresses nonspecific signals and improves the detection limit. (A)Schematic of workflow modifications to reduce nonspecific puncta: protease treatment to remove membrane-associated RNP aggregates and transient glutamine deprivation to reduce intranuclear nonspecific accumulation. (B)Representative z-projection image of an OPM volume acquired from MAC-positive HT1080 cells (acquisition time, ~1.5 s per volume). Nuclei and CRISPR spots are shown in magenta and green, respectively. Scale bar, 100 μm. (C)Representative intranuclear CRISPR spots. Maximum-intensity z-projections of the GFP channel are shown; nuclear boundaries are outlined in yellow and arrows indicate discrete spots. Scale bar, 5 µm. (D)Cell-level SNR distributions without and with the workflow optimizations. Vertical lines denote the SNR threshold used for calling and the corresponding limit of detection (LoD) under the defined criteria. Blue, positive-control cells (n = 16,831 without optimization; n = 13,056 with optimization); orange, negative-control cells (n = 20,127 without optimization; n = 16,994 with optimization). (E)Precision–recall curves without and with the workflow optimizations by sweeping the SNR threshold on datasets from 30,723 MAC-positive cells and 37,577 MAC-negative cells across three independent experiments. mAP, mean average precision (without optimization, 0.941; with optimization, 0.971) and standard deviation (shadow).

To improve performance at lower target frequencies, we focused on reducing cell-level false positives, which directly constrain the SNR threshold required to satisfy the detection-limit criteria. We therefore refined the labeling workflow to address two practical sources of spurious signals: intranuclear nonspecific puncta and membrane-associated RNP aggregates (Fig. 3A).

First, motivated by reports that nuclear bodies can concentrate proteins through liquid– liquid phase separation (*21*), we tested whether transient nutrient deprivation after electroporation would reduce nonspecific intranuclear accumulation. HT1080 cells were deprived of fetal bovine serum, glutamine, or glucose for 8 h after electroporation and imaged at the end of the deprivation period. Both glutamine and glucose deprivation reduced nonspecific calls in negative-control cells (Fig. S4), but only glutamine deprivation was therefore used in subsequent experiments since glucose deprivation impaired cell adherence. Varying the deprivation duration further reduced nonspecific calls and reached a plateau after 24 h (Fig. S5). Second, we introduced brief protease treatment (Trypsin or Accumax) to remove membrane aggregates that otherwise became internalized and generated false-positive puncta (Fig. S6). By integrating metabolic modulation and protease treatment into the labeling workflow, we reduced nonspecific fluorescence on cell membranes (Fig. 3B) and decreased intranuclear spurious puncta within nuclei (Fig. 3C). Consistent with these improvements, the false-positive population previously visible in the per-cell SNR histograms was largely eliminated, enabling lower SNR thresholds to meet the target detection limits (Fig. 3D). Under the defined imaging and analysis pipeline, the Hi-CRI LoD improved from approximately 0.1% to 0.01%, with corresponding gains in precision-recall performance and mAP (mAP, 0.941 to 0.971) relative to the unoptimized workflow (Fig. 3E).

### Application to MMCT reveals substantial pre-readout attrition

To test whether Hi-CRI can quantify low-frequency chromosome transfer events, we applied it to microcell-mediated chromosome transfer (MMCT) of a mouse artificial chromosome (MAC) into HT1080 recipient cells (Fig. 4A). After microcell fusion and a brief recovery period (1 day), we performed CRISPR-based imaging and screened large populations by high-throughput OPM acquisition and analysis (Fig. 4B). To reduce extranuclear artifacts, cell calling was restricted to intranuclear signals within the segmented nucleus and excluded extranuclear puncta.

**Fig. 4.**
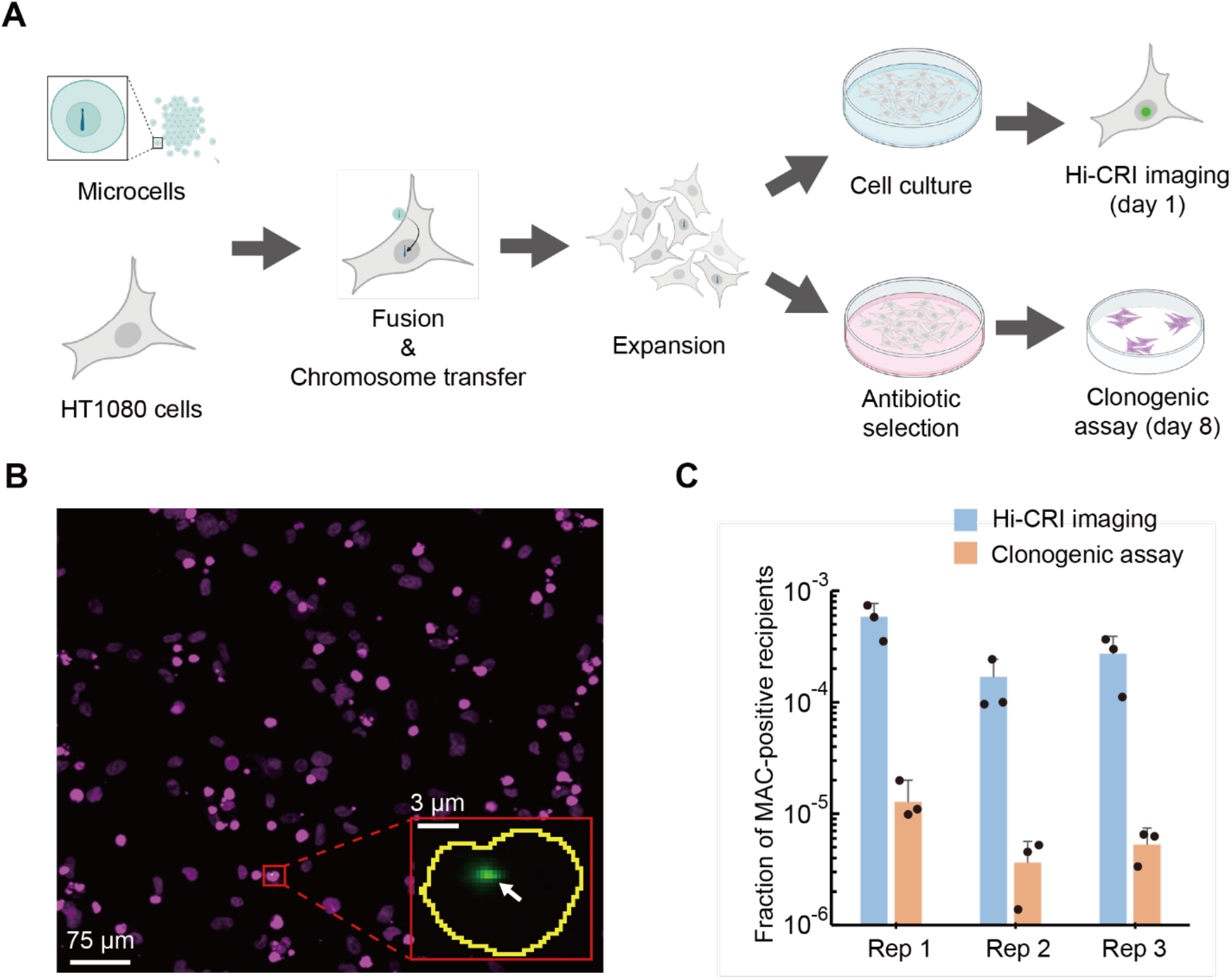
Quantifying low-frequency MAC-positive recipients after MMCT using Hi-CRI. (A) Experimental schematic. Following microcell fusion and expansion, recipient HT1080 cells were split into two arms: Hi-CRI imaging after 1 day of culture, or antibiotic selection followed by a clonogenic assay after 8 days. (B) Representative z-projection from an OPM volume in the post-MMCT population. Nuclei (magenta) and CRISPR spots (green) are shown. The inset shows a MAC-positive cell, in which the nuclear boundary is outlined in yellow and the arrow marks an intranuclear CRISPR spot. Scale bars, 75 µm (overview) and 3 µm (inset). (C) Estimated MAC-positive fraction measured by Hi-CRI imaging and by antibiotic-selection-based clonogenic assay across three independent MMCT experiments (Rep 1–3; y-axis, log scale). For Hi-CRI, the fraction is the number of cells called positive divided by the number of analyzed cells; for the clonogenic assay, the fraction is the number of surviving colonies divided by the number of input cells. Dots indicate technical replicates, ranging from 14,097 to 33,032 cells per replicate; error bars indicate ± s.d.

Across three independent MMCT experiments, Hi-CRI analyzed 184,235 recipient cells in total (~60,000 cells per experiment) and estimated a 0.03% MAC-positive fraction (56 events; 95% confidence interval, 0.022 to 0.038%; Fig. 4C). In parallel, an antibiotic-selection-based clonogenic assay performed on the same batch yielded a markedly lower MAC-positive estimate (0.0007%, Fig. 4C). This discrepancy, measured at different post-fusion readout stages, is consistent with substantial pre-readout attrition during antibiotic selection and clonogenic outgrowth and suggests that only a small fraction of early MAC-positive cells ultimately form antibiotic-resistant colonies. Together, these results establish Hi-CRI as a selection-independent readout for process-level evaluation of chromosome transfer.

## Discussion

This study establishes Hi-CRI as an integrated platform for quantifying low-frequency chromosomal events in living cells by combining engineered RNP-based labeling, workflow-level suppression of nonspecific aggregates, rapid OPM-based volumetric imaging, GPU-accelerated image analysis, and an explicit detection-limit framework. After workflow optimization, Hi-CRI achieved a detection limit of 0.01% for the target-positive cell fraction under the α/β criteria used here.

In an MMCT setting, Hi-CRI measured a 0.03% MAC-positive fraction among 184,235 recipients (95% confidence interval, 0.022 to 0.038%), substantially higher than the estimate obtained by an antibiotic-selection-based clonogenic assay (0.0007%). Because Hi-CRI captures early nuclear-localized transfer events while the clonogenic assay requires long-term survival under selection, the discrepancy is consistent with substantial pre-readout attrition between these stages.

A central advance of Hi-CRI is that sensitivity claims in CRISPR imaging are explicitly tied to an error-controlled detection-limit definition. Rare-event measurements are interpretable only when false positives and false negatives are specified at the same decision level as the reported frequency. By implementing cell-level SNR-based calling within an IUPAC-aligned α/β framework, Hi-CRI enables experiment-size-aware control of false positives and the expected number of false discoveries. Importantly, this formalization also turns workflow development into a quantitative optimization problem: by evaluating detection limits on large control cohorts, we could identify nonspecific aggregates as the dominant source of cell-level false positives and systematically refine labeling conditions. Guided by these criteria, workflow-level suppression of nonspecific aggregates improved the detection limit from ~0.1% to 0.01% under the defined acquisition and calling settings. This provides a transferable recipe for reporting detection limits in CRISPR imaging and enables meaningful comparisons across instruments, targets, and analysis pipelines.

The MMCT results highlight the distinction between early physical presence of a transferred chromosome and long-term functional establishment under selection. Because Hi-CRI restricts calling to intranuclear signals within segmented primary nuclei, early positives are consistent with nuclear-localized MAC signals rather than extranuclear artifacts. The subsequent reduction in clonogenic readout is therefore unlikely to arise from events occurring entirely outside the nucleus but suggests that many early MAC-positive cells fail to progress through post-entry bottlenecks required for stable outgrowth. Potential contributors include incomplete stabilization within the recipient nuclear environment (*22*–*26*), fusion-associated stress responses (*27*), limited compatibility with host cell-cycle progression (*28, 29*), insufficient chromosomal maturation, or impaired chromosome segregation during subsequent mitotic divisions, any of which could limit progression to stable clonogenic expansion. Notably, cell-cycle compatibility has been shown to influence nuclear stability and chromosomal integrity in related contexts such as somatic cell nuclear transfer and cell fusion (*28, 29*), supporting the plausibility of such post-nuclear constraints. These mechanisms are not directly tested here, and longitudinal studies that track early Hi-CRI positives through subsequent culture and selection will be required to link early nuclear detection to long-term establishment.

Several limitations remain. First, analysis throughput lags behind imaging throughput in the current single-GPU configuration, motivating multi-GPU scaling and on-the-fly inference. Second, detection at small, low-copy loci will require improved effective SNR, for example through optimized guide tiling, brighter or switchable reporters, and further background suppression. Third, while Hi-CRI identifies positives at high throughput, methods for retrieving or sorting those cells remain to be developed. Finally, broader generality will benefit from benchmarking across additional cell types, endogenous loci, and instrument configurations under clearly specified acquisition and calling criteria.

In chromosome engineering workflows, including MMCT, Hi-CRI provides a viability-preserving, selection-independent readout that quantifies the “silent majority” of early chromosome-transfer events that can inform design–build–test–learn cycles. More broadly, the platform supports studies of rare karyotypic states in heterogeneous systems, including longitudinal tracking of subclones under perturbation. By linking live-cell genomic detection to an explicit detection-limit control, Hi-CRI establishes a quantitative and statistically interpretable framework for decision-making in settings where rare chromosomal states are consequential.

## Materials and Methods

### DNA target selection

DNA targets on human chromosome 13, 19 and the major satellite region of mouse chromosomes were designed based on genomic sequence or selected from previous studies (*10, 12, 30*). 12-nt of specific sequences are incorporated into the sgRNAs for the CRISPR-based DNA labeling as shown in Tables S2.

### Plasmid construction and preparation

To construct the plasmid for dCas9 expression and purification, pET-Hisx6-NLS-SpCas9-NLS (a gift from the Nishimasu lab at UTokyo) was modified with D10A/H840A point mutations on the SpCas9 sequence. For the basic NLS configurations, NLSs, as shown in Tables S1, were fused at the N-terminus of dCas9. In the NLS combination, 2xNLS from NLP, SV40, and c-Myc were fused at the N-terminus, the middle, and the C-terminus of the dCas9-GFP fusion, respectively. The GFP variants (sfGFP, GenBank: ASL68970; mNeonGreen, GenBank: AGG56535; mWasabi, GenBank: ABW74902.1; copGFP, GenBank: AAQ01184) were fused at the C-terminus of dCas9, followed by an N-terminal 6xHis tag. To construct the dCas9–RFP plasmid, the GFP fusion in the previous construct was replaced with a 2x mScarlet fusion (GenBank: APD76535).

For in vitro T7 sgRNA transcription, pET-sgRNA incorporating the synthesized sgRNA scaffold (IDT), as shown in Tables S2, was constructed as the DNA template for PCR amplification.

All plasmids were prepared using NEB 10-beta Competent E. coli (High Efficiency) (NEB, C3019H) and purified with the Monarch Plasmid Miniprep Kit (NEB, #T1010). The plasmids were quantified using a NanoDrop spectrophotometer (Thermo Fisher Scientific).

### Protein expression and purification in vitro

The expression and purification of dCas9-GFP proteins were performed according to the protocol from the Nishimasu lab at UTokyo (*31*). Briefly, expression plasmids were transformed into Rosetta 2 (DE3) competent cells (Novagen, #71397-4CN). The transformed E. coli was cultured in LB medium (BD, #244610) with 50 µg/ml kanamycin (Wako, #117-00961) at 37 °C until reaching an OD600 of 0.8. After a 20-minute cold shock on ice, isopropyl β-D-1-thiogalactopyranoside (IPTG;Wako, #094-05323) was added to a final concentration of 0.1 mM. The cells were then harvested by centrifugation (10,000 x g, 3 min) after culturing at 20 °C for 20 hours. The resulting cell pellets were either stored at −80 °C or immediately used for protein purification.

The cell pellets were resuspended in the lysis buffer (20 mM Tris-HCl pH 8.0, 1 M NaCl, 20 mM imidazole) and sonicated on ice. The lysates were centrifuged at 10,000 x g for 10 minutes at 4 °C to pellet the cellular debris. Proteins in the supernatant were purified using a column (Bio-Rad, #7311550) filled with Ni-NTA agarose (QIAGEN, #30210). The supernatant was loaded onto the column, which had been pre-equilibrated with the lysis buffer. After the supernatant passed through, the column was sequentially washed with the lysis buffer followed by the washing buffer (20 mM Tris-HCl pH 8.0, 0.3 M NaCl, 20 mM imidazole). The elution buffer (20 mM Tris-HCl pH 8.0, 0.3 M NaCl, 300 mM imidazole) was then applied to the column to elute the protein. The collected flow-through was further purified using the column filled with SP Sepharose High Performance (Cytiva, #17108701), which was washed and pre-equilibrated with Buffer A (20 mM Tris-HCl pH 8.0, 0.25 M NaCl). After loading the sample, the column was washed with Buffer A, and the protein was eluted using Buffer B (20 mM Tris-HCl pH 8.0, 1 M NaCl). The eluted protein was then concentrated using Amicon Ultra Centrifugal Filters (100 kDa MWCO; Millipore, #UFC510008), quantified by NanoDrop (Thermo Fisher Scientific), and divided into aliquots. Each aliquot was flash-frozen in liquid nitrogen and stored at −80 °C.

### *In vitro* sgRNA transcription

sgRNAs were transcribed in vitro using the HiScribe T7 Quick High Yield RNA Synthesis Kit (NEB, #E2050S). PCR templates were amplified from pET-sgRNA with oligo primers (IDT) as shown in Tables S3 using Q5 High-Fidelity DNA Polymerase (NEB, #M0491). The sgRNAs were purified using the Monarch RNA Cleanup Kit (50 µg, NEB, #T2047L), analyzed by gel electrophoresis, and quantified by NanoDrop (Thermo Fisher Scientific). Aliquots of sgRNAs were flash-frozen in liquid nitrogen and stored at −80 °C.

### Cell culture

U2OS and HT1080 cells were cultured in D-MEM (High Glucose) with L-Glutamine and Phenol Red (Wako, #044-29765), supplemented with 10% FBS (Sigma, #F7524) and 1% Antibiotic-Antimycotic (100X) (Thermo Fisher Scientific, #15240096). For the MAC-positive HT1080 cell line (HT1080 MI-MAC27-2), G418 (InvivoGen, #ant-gn-1) was added to the medium at 600 µg/ml. CHO-K1 cells were cultured in Ham’s F-12 with L-Glutamine and Phenol Red (Wako, #087-08335), supplemented with 10% FBS and 1% Antibiotic-Antimycotic. For the MAC-positive CHO cell line (CHO-K1 MI-MAC2), Hygromycin (Wako, #085-06153) was added at 300 µg/ml. K562 and Jurkat cells were cultured in RPMI-1640 with L-Glutamine and Phenol Red (Wako, #189-02025), supplemented with 10% FBS and 1% Antibiotic-Antimycotic. All cells were maintained at 37°C with 5% CO_2_ in a humidified incubator.

U2OS cells were provided by the Miyaoka lab (Tokyo Metropolitan Institute of Medical Science). HT1080 and CHO-K1 cells were obtained from Yasuhiro Kazuki’s lab (Tottori University). K562 (#RCB0027) and Jurkat (#RCB3052) cells were provided by RIKEN BRC.

### MV-MMCT

For microcell preparation, CHO cells retaining MAC (MI-MAC2) (*32*) were used as donor cells, and microcells were obtained from them. The MMCT for microcell formation used the MV-MMCT method reported in prior research (*33*). For recipient cell preparation, cells were seeded in a 100 mm dish for 96 hours before fusion. In the day prior to fusion, 3 × 10^6 cells were seeded in each 60 mm dish. On the day of fusion, cryopreserved microcells, obtained from Yasuhiro Kazuki’s lab (Tottori University) *(8, 9)*, were incubated at 37 °C for 1 minute in a water bath. The microcells were then resuspended in 10 mL of serum-free DMEM and centrifuged at 2,000 rpm for 10 minutes. This washing step was repeated once more. Finally, the microcells were resuspended in 3-4 mL DMEM supplemented with 10% FBS for co-culturing with recipient cells for 24 hours.

### Clonogenic assay

The cells were seeded in five 100 mm dishes. 300 µg/ml Hygromycin or 600 µg/ml G418 was added into the culture media for drug selection for 8 days. Drug-resistant colonies were fixed in methanol for 5 minutes. The fixed samples were incubated in 5% Giemsa staining solution (MERCK, Darmstadt) for 15 minutes. After staining, the samples were washed with water, and stained colonies were counted.

### Nutrition deprivation

HT1080 cells were subjected to nutrient deprivation by culturing in the corresponding D-MEM medium: For Gln deprivation, D-MEM without L-glutamine (Wako, #040-30095) supplemented with 10% FBS (Sigma, #F7524) and 1% Antibiotic-Antimycotic (100X) (Thermo Fisher Scientific, #15240096); For Glu deprivation, D-MEM without glucose (Wako, #042-32255) supplemented with 10% FBS (Sigma, #F7524) and 1% Antibiotic-Antimycotic (100X) (Thermo Fisher Scientific, #15240096); For FBS deprivation, D-MEM (Wako, #044-32955) supplemented with 1% Antibiotic-Antimycotic (100X) (Thermo Fisher Scientific, #15240096).

MAC-negative cells were starved for 8 h in the indicated deprivation medium and then analyzed by the Hi-CRI workflow. To determine the time-dependent effect of glutamine deprivation, cells were cultured under pre-RNP-delivery and post-RNP-delivery Gln-deprivation regimens. Post-RNP-delivery Gln deprivation was fixed at 8 h, whereas pre-RNP-delivery Gln deprivation was varied to extend the total deprivation time. In the optimized protocol, cells are cultured under pre-RNP-delivery Gln deprivation for 16 h and under post-RNP-delivery Gln deprivation for 8 h, followed by imaging.

### Protease treatment

Cells were washed with PBS, centrifuged at 200 g for 3 minutes and resuspended in 100 µl of either Trypsin-EDTA (0.25%) (Thermo Fisher Scientific, #25200056) or Accumax (Innovative Cell Technologies, #AM105). Cells were incubated at 37°C for 3 minutes with Trypsin-EDTA or at room temperature for 10 minutes with Accumax. To remove RNP aggregates on cell membranes, cells were first treated with Accumax, followed by PBS washing. Subsequently, cells were incubated with Trypsin-EDTA and washed again with culture medium.

### RNP delivery

RNPs were delivered into the cells via electroporation using the Amaxa 4D-Nucleofector (Lonza). For U2OS cells (program CM-104) and Jurkat cells (program CL-120), the SE kit (Lonza, #V4XC-1032) was used. For K562 cells (program FF-120), CHO-K1 cells (program DT-133), and HT1080 cells (program FF-113), the SF kit (Lonza, #V4XC-2032) was used.

Briefly, 2.5 × 10^5 cells were harvested, washed with PBS, and resuspended in a 15 ul EP buffer before electroporation. For RNP assembly *in vitro*, 25 pmol dCas9-2xGFP was pre-mixed with 100 pmol sgRNAs in a 9 µL solution at room temperature for 10 minutes. RNPs and cells were mixed in the 20 µl Nucleocuvette Strip with 1 µl of 100 µM Alt-R Cas9 Electroporation Enhancer (IDT, #1075916) and 0.5 µl of 50% glycerol. Following electroporation, the Nucleocuvettes were incubated at room temperature for 10 minutes, after which the cells were washed with PBS and subjected to protease treatment. Subsequently, cells were passed through a 20 µm cell strainer (pluriSelect, #43-10020-40) and seeded into each well of a 96-well glass-bottom plate (ibidi, #89607) pretreated with 3 mg/ml collagen (Nitta Gelatin Inc, #631-00771).

### Confocal and light-sheet microscopy

Confocal imaging was conducted using the Nikon A1R confocal microscope, which is equipped with a CCD camera and a 60X, NA-1.4 oil immersion objective. Hoechst 33342 (Sigma, #875756-97-1) was excited with a 405 nm laser, and its emission was collected through a 450/50 nm filter. GFP was excited using a 488 nm laser, with its emission collected through a 525/50 nm filter. Stack imaging was performed with a z-axis step size of 0.17 µm and a scanning speed of 0.5 frames per second.

For light-sheet microscopy, the design of the single-objective light-sheet microscope is described in a previous study *(15–17)*. In this study, a 20X, NA-0.75 dry objective (Olympus) was used. Stack imaging was performed with a camera frame rate of 300 Hz and a light-sheet scanning speed of 0.25 mm/s. A 488 nm laser was employed to excite GFPs, while a 637 nm laser was used to excite NucSpot Live 650 (Biotium, #40082-T).

Cell nuclei were stained in culture media with 2.5 µg/ml Hoechst 33342 for 10 minutes or 0.5x NucSpot Live 650 for 2 hours before imaging. Live cells were maintained in a 37°C chamber during imaging.

### Image processing

Images obtained from confocal microscopy were processed using Fiji (ImageJ). For presentation, Z-projections were generated from the image stack and cropped, followed by brightness and contrast adjustments in Fiji.

Images obtained from OPM were reconstructed and processed using JupyterLab. The source codes are listed at Table S4. The source of raw image data is provided in Table S5. Briefly, image shift between green and far-red channels was calculated using images of multicolor beads after 3D reconstruction. The calculated shift parameters in the x, y, and z dimensions were used to compensate for the shift among different channels. The 3D views were generated using Napari, with adjustments to the minimum and maximum values.

### Signal-to-noise ratio (SNR) analysis

For images obtained from confocal microscopy, the Hoechst channel was z-maximum projected and used to generate single-nucleus masks. These masks were applied to the corresponding GFP images. Within each nucleus, candidate spots were identified as local intensity maxima in the GFP channel, and the peak pixel intensity for each focus was recorded. To estimate noise, pixels belonging to putative spots were excluded (by masking around detected peaks), and the standard deviation of the remaining nuclear pixels was computed.The SNR was calculated by dividing the peak fluorescence intensity of the spots by the standard deviation of noise within the nucleus.

For images obtained from APOM, in the GFP channel, Voronoi-Otsu labeling and a size filter are applied to exclude the largest 5% of objects. Subsequently, 3D masks, generated via 3D segmentation in the far-red channel for each nucleus, along with a gradient-based spot detection algorithm, are applied to the GFP channel to identify CRISPR spots within the nucleus. The SNR is calculated as described above.

### Relative fluorescence ratio calculation

For single-cell analysis, fluorescence intensity profiles were extracted across the nucleus along manually defined line segments passing through the nuclear center. Hoechst images were used to delineate the nuclear boundary, and the corresponding GFP channel was sampled along the same line. The cytoplasmic regions were defined as flanking segments on both sides of the nucleus, excluding the nuclear boundary by 1–2 µm to avoid edge artifacts. Mean GFP intensity values were measured separately in the nuclear and cytoplasmic regions of each line profile. The RFR was then calculated as the mean nuclear intensity divided by the mean cytoplasmic intensity.

### Intersection-over-union (IoU) evaluation

To quantify the colocalization between GFP and RFP signals, the intersection-over-union (IoU) coefficient was calculated for each detected focus. Binary masks for GFP and RFP spots were generated from z-projected images by intensity thresholding followed by morphological filtering to remove small background pixels. For each pair of masks, the intersection area and the union area were measured, and IoU was defined as the intersection area divided by the union area. A focus pair was considered colocalized when IoU ≥ 0.5.

### Precision–recall (PR) analysis

Precision was calculated as the ratio of true positives to the sum of true positives and false positives, while recall was determined as the ratio of true positives to the sum of true positives and false negatives. Precision and recall curves were generated to evaluate the detection performance. For each SNR threshold value, precision and recall were computed, and the curve was plotted by decreasing SNR threshold across the entire range.

### Limit of detection (LoD) evaluation

LoD was evaluated following the International Union of Pure and Applied Chemistry (IUPAC) framework using separate positive and negative reference populations. For each experiment, Hi-CRI data were acquired from a COI-positive cell line (all cells harboring the chromosome of interest) and a COI-negative cell line (no COI). For every nucleus, the SNR of each CRISPR focus was computed as described above, and the brightest focus per nucleus was used as the decision variable.

To construct error profiles, an SNR decision threshold *T* was swept across the observed SNR range. At each threshold, nuclei were classified as “detected” if at least one focus had SNR ≥ *T*; otherwise, they were classified as “not detected.” The false-positive rate (FPR) was calculated as the fraction of COI-negative nuclei classified as detected, and the false-negative rate (FNR) as the fraction of COI-positive nuclei classified as not detected. Operational LoD values were then derived by identifying SNR thresholds that satisfied the IUPAC-style criteria, i.e., an upper bound on FPR (type I error) and FNR < 0.5 (type II error), treating the fraction of COI-positive cells as the analyte level.

### Statistical analysis

The significance of differences was assessed through the following steps: 1) Evaluation of the homogeneity of variances using Levene’s and Bartlett’s tests; 2) Calculation of p-values using two-sided tests.

For cases with homogeneous variances, independent samples t-tests were employed to compare two groups, followed by Tukey’s Honestly Significant Difference (HSD) test for post hoc analysis. In instances of non-homogeneous variances, Welch’s t-test was applied for comparing two groups.

## Supporting information

Supplemental Infomation

## Acknowledgments

This study was performed in part at the Tottori Bio Frontier managed by Tottori prefecture.

## Author contributions

Conceptualization: HX, SO, KY, YK

Methodology: HX, YI, KY, YM

Investigation: HX, NK

Software: YI, HX

Formal analysis: YI, HX

Visualization: HX, YI

Writing—original draft: HX, YI, SO

Writing—review &

editing: HX, YI, KY, NK, NO, HM, YK, SO

## Data and materials availability

All code and raw data associated with this manuscript is hosted on Zenodo, listed in Table S4 and Table S5.

