## Supplemental Infomation for "High-throughput CRISPR live-cell imaging of low-frequency chromosomal events quantifies the latent efficiency of chromosome engineering"

**Supplementary Materials**

Figs. S1 to S6

Tables S1 to S5

Data S1

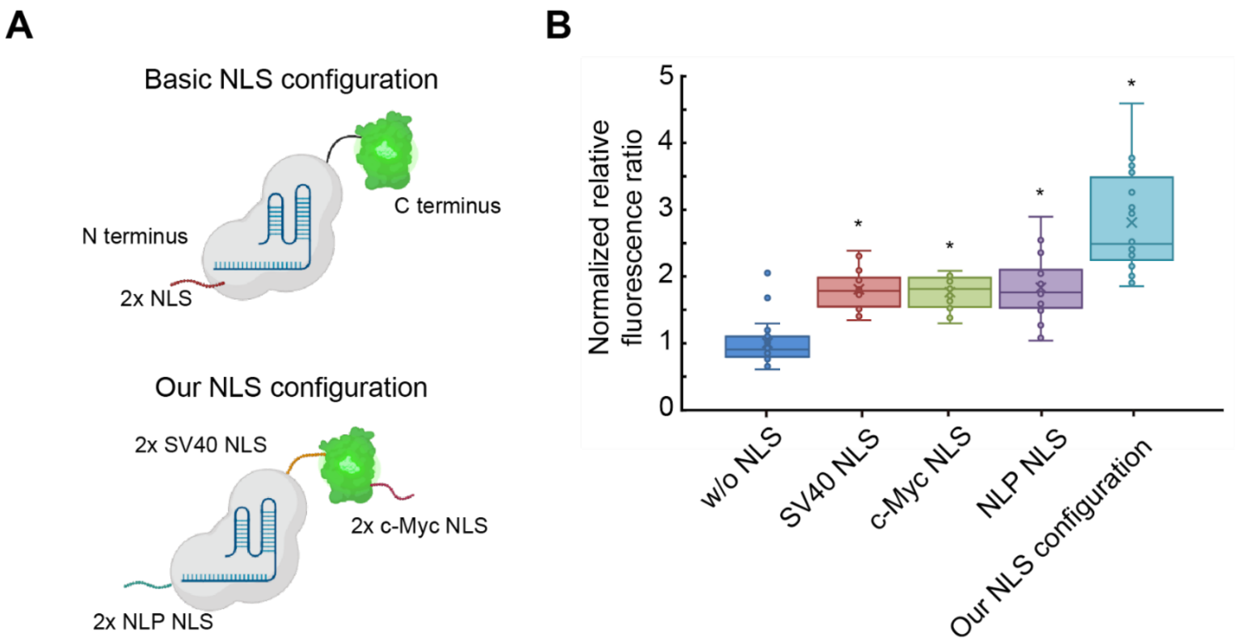


**Fig. S1. Screening nuclear localization signals (NLSs) to enhance dCas9 RNP nuclear accumulation.**

**(A)** The NLS configuration of dCas9 RNPs.

**(B)** Normalized relative fluorescence ratio between the nucleus and cytoplasm. 3 independent experiments, $n > 14$. * $p < 0.05$ vs w/o NLS (two-sided Student's t-test, pairwise comparisons).

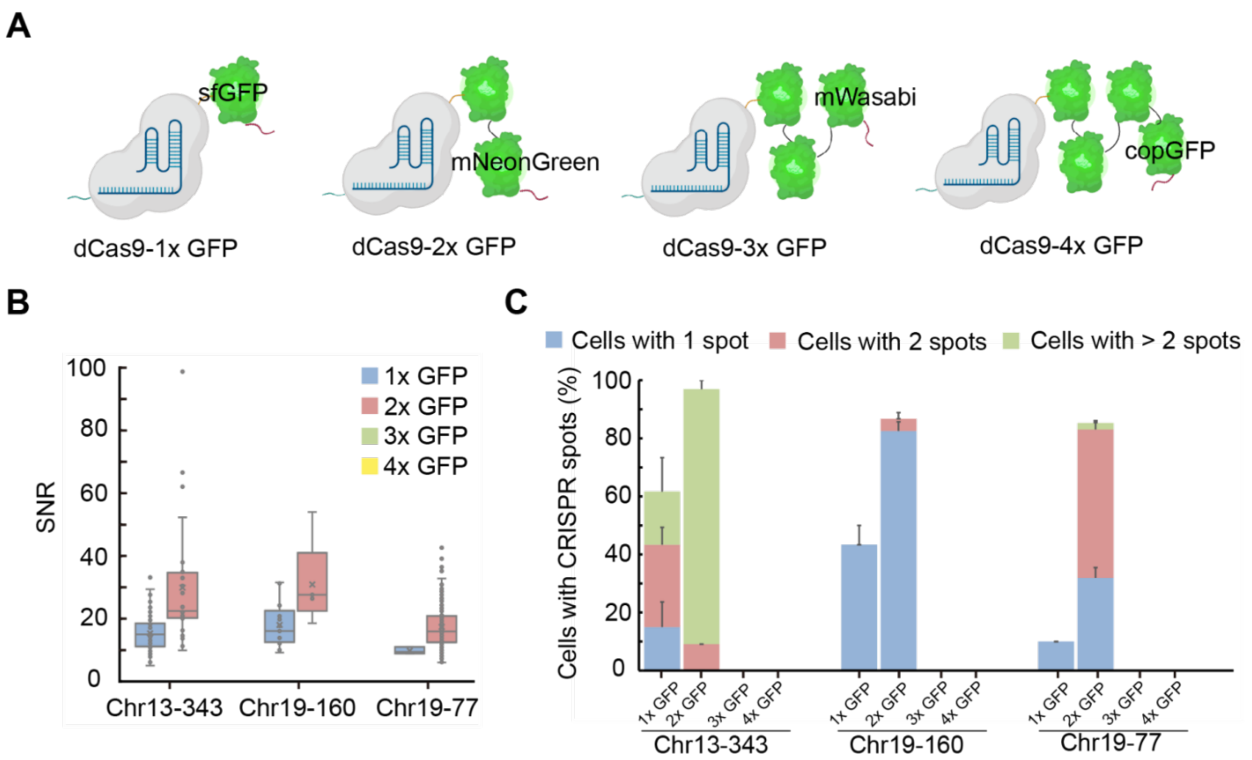


**Fig. S2. Optimization of GFP fusions with dCas9 RNPs to enhance chromosome labeling.**

**(A)** Schematic of GFP fusions with dCas9 RNP

**(B)** SNR of CRISPR signals. Target labels are shown below; for example, "chr13" denotes human chromosome 13 and "343" indicates the copy number (repeat count) of the target sequence. 3xGFP and 4xGFP fusions failed to generate detectable CRISPR signals. Three independent experiments, n > 30 cells.

**(C)** The percentage of cells containing CRISPR signals. 3xGFP and 4xGFP fusions failed to generate detectable CRISPR signals. 3 independent experiments, n > 30 cells. Data shows mean ± s.d.

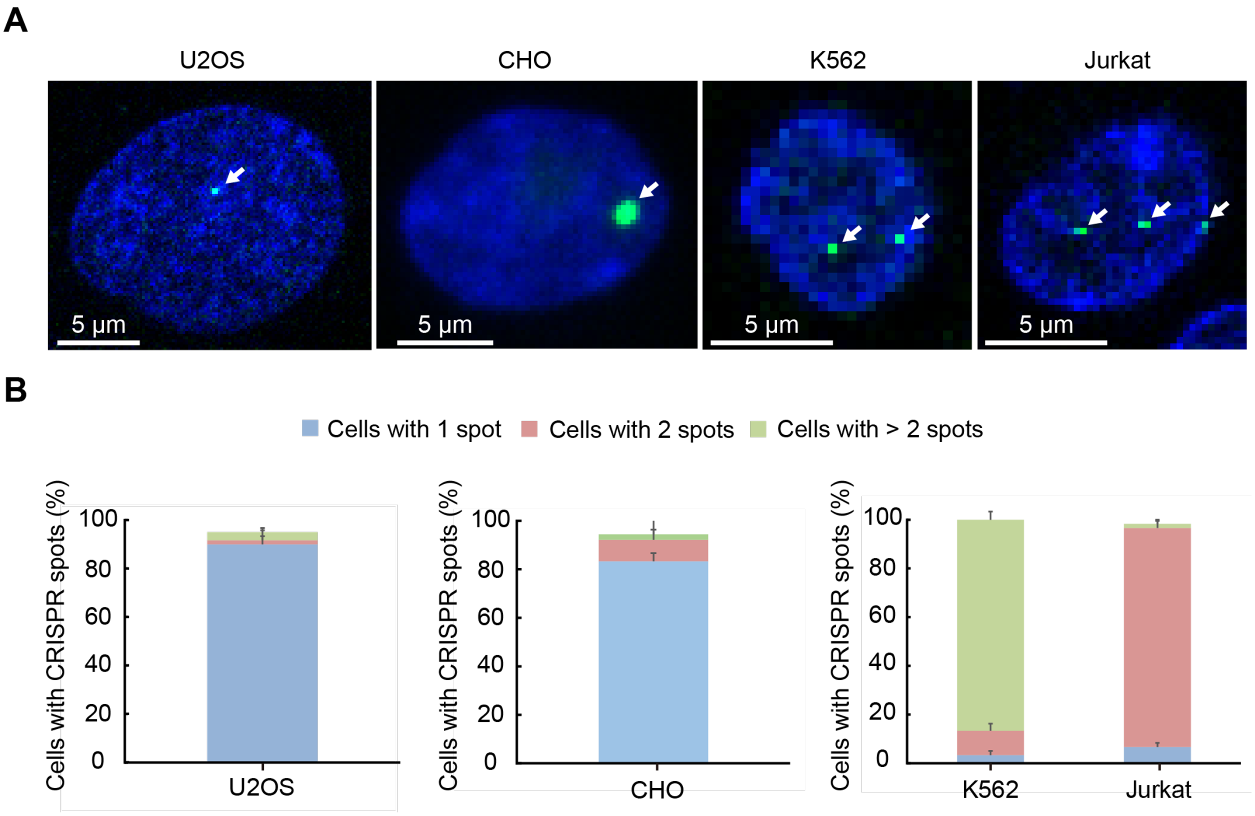


**Fig. S3. Live-cell chromosome labeling across multiple cell types.**

**(A)** Representative z-projection images of live-cell chromosome labeling across multiple cell lines. Human chromosome 19 (Chr19) is labeled in U2OS, K562, and Jurkat cells, and the mouse artificial chromosome (MAC) is labeled in a MAC-positive CHO cell line. Arrows indicate CRISPR spots. Scale bars, 5 µm.

**(B)** Fraction of cells exhibiting CRISPR spots. 3 independent experiments, n > 30 cells. Data shows mean ± s.d.

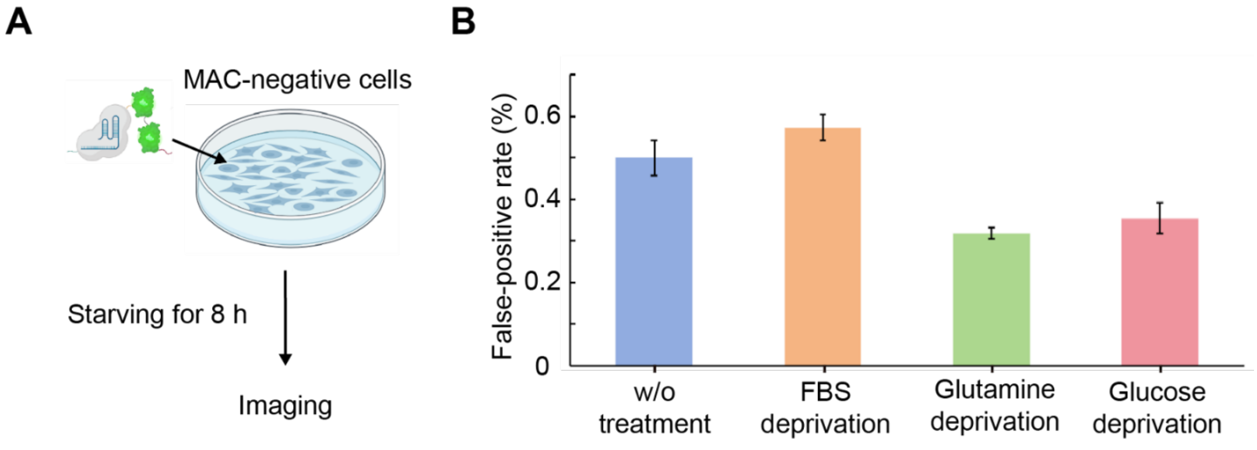


**Fig. S4. Effect of nutrient deprivation on the false-positive rate.**

**(A)** Schematics of nutrition deprivation. MAC-negative cells were incubated for 8 h under deprivations of FBS, glutamine, or glucose, and then analyzed by the Hi-CRI workflow. Cells in the control groups (w/o treatment) incubated under no deprivation.

**(B)** False-positive rates (FPR, %) under nutrient deprivations. All samples were analyzed using a fixed SNR threshold of 20. Three replicates, n = 30,000 cells; Bars show mean ± s.d.

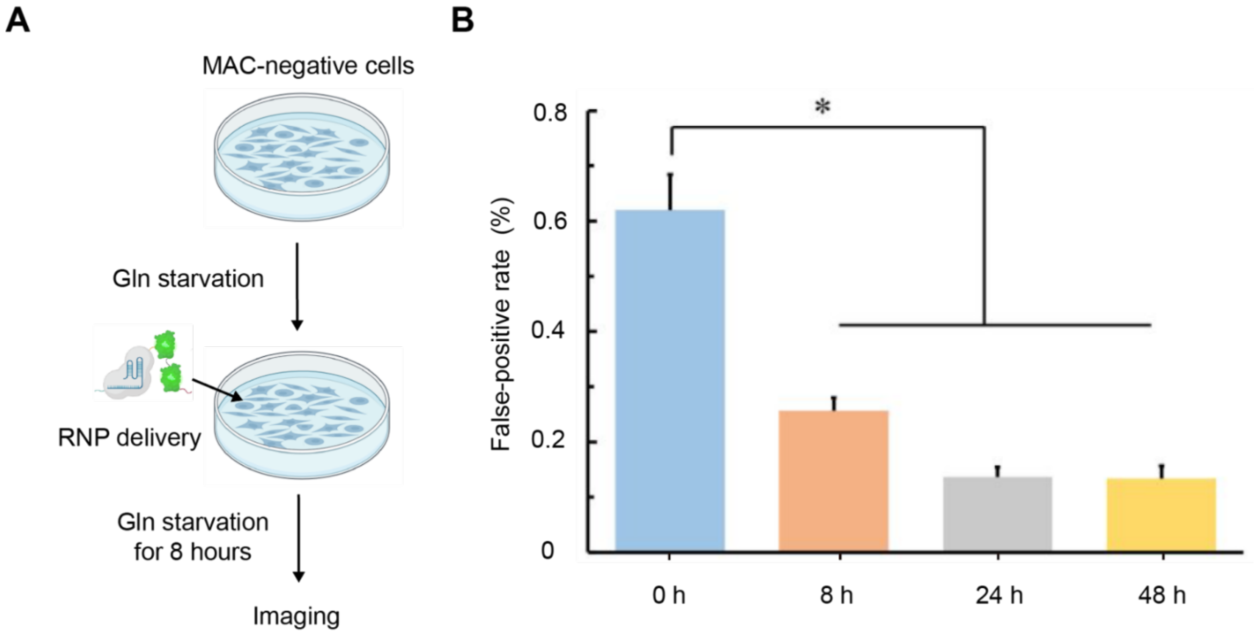


**Fig. S5. Time-dependent effect of glutamine deprivation on the false-positive rate.**

**(A)** Schematic of the glutamine-deprivation regimen. The second glutamine starvation was fixed at 8 hours, while the first was adjusted according to the total starvation time.

**(B)** False-positive rate (FPR) after 0, 8, 24, or 48 h of deprivation. Cells were starved before and after RNP delivery; quantification used a fixed SNR threshold of 20. Three replicates, n = 30,000 cells; * $p < 0.05$ vs 0 h (two-sided Student's t-test, pairwise comparisons); Bars, mean ± s.d.

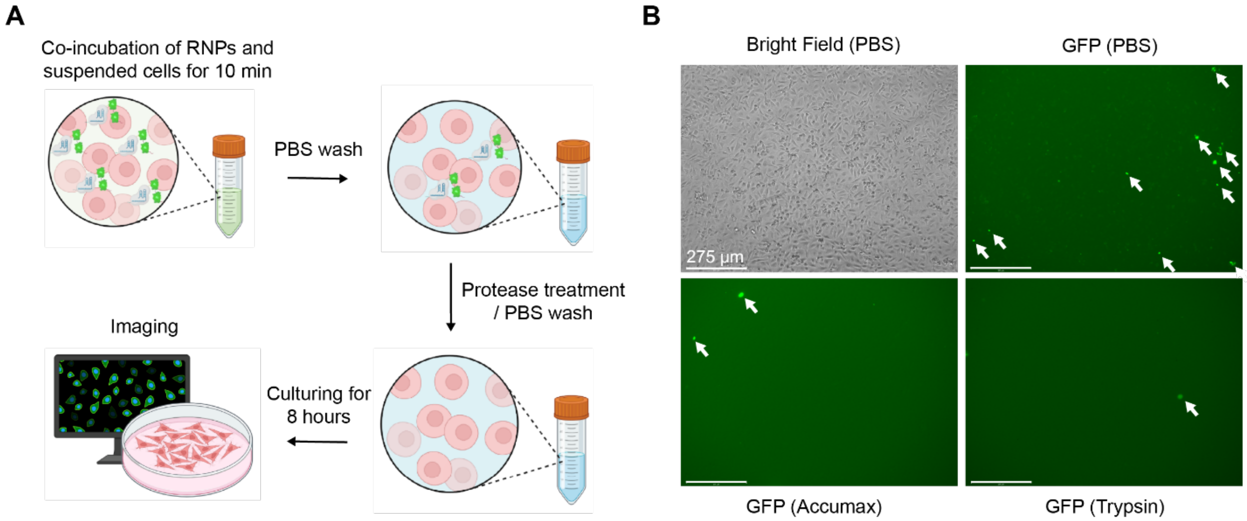


**Fig. S6. Removal of extracellular GFP fluorescence.**

**(A)** Workflow schematic. Suspended cells were co-incubated with dCas9–GFP RNPs for 10 min, followed by sequential PBS washes and optional protease treatments (Accumax or trypsin or PBS) to eliminate surface-bound complexes. Cells were then replated and cultured for 8 h before imaging.

**(B)** Representative images. Control (PBS) shows cells without protease treatment. Arrows indicate GFP aggregations. Scale bars, 275 µm

**Table S1. Sequence of NLSs used in this study**

| NLS name | AA sequence (N-terminus to C-terminus) | DNA sequence (5’ to 3’) |
|---|---|---|
| SV40 | PKKKRKV | ccaaagaagaagcggaaggtc |
| c-Myc | PAAKRVKLD | ccggctgcgaaacgggttaaactcgat |
| NLP NLS | KRPAATKKAGQAKKKK | aaaaggccggcggccacgaaaaaggccggccaggcaaaaaagaaaaag |

**Table S2. Sequences of sgRNAs used in this study**

sgRNA scaffold (the spacer is underlined):

5’-NNNNNNNNNNNNguuuaagagcuaugcuggaaacagcauagcaaguuuaaauaaggcuaguccguuaucaacuugaaaaaguggcaccgagucggugcuuuuuuu-3’

| Target name | Target region | Spacer sequence | Copy number of target |
|---|---|---|---|
| Chr13-343 | Chr13q34 | GGACCATTCCTTC | 343 |
| Chr19-160 | Chr19p12 | GTGACAGTGAAC | 160 |
| Chr19-77 | Chr19q13.43 | GAGGAGGGAAGC | 77 |
| MajSat_1 | Mouse major satellite | GGACGTGGAATA | \ |
| MajSat_2 | Mouse major satellite | GGACCTGGAAT | \ |
| Nonsense | \ | GGAGTTGTGTTTGTGGACGAAG | \ |

**Table S3. Co-localization rate**

| sgRNAs complexed with dCas9–GFP | sgRNAs complexed with dCas9–RFP | Mean of co-localization rate (%) | STD of co-localization rate (%) |
|---|---|---|---|
| sgMajSat_1 | sgNonsense | 2.3 | 2.4 |
| sgNonsense | sgMajSat_2 | 0.9 | 0.1 |
| sgMajSat_1 | sgMajSat_2 | 96.0 | 0.5 |

**Table S4. List of analysis scripts used in this study**

| File name | Function | Repository |
|---|---|---|
| 3D reconstruction | 3D reconstruction | 10.5281/zenodo.17596985 |
| step1_shift_calculation | Image analysis | |
| step2_spot_identifiction_v3 | Image analysis | |
| step3_analysis_of result_v3 | Image analysis | |

**Table S5. Raw 3D image datasets used in this study**

| Figure | Repository |
|---|---|
| Fig. 2 | 10.5281/zenodo.17599633 |
| Fig. 3 | 10.5281/zenodo.15240624<br>10.5281/zenodo.15241187 |
| Fig. 4 | 10.5281/zenodo.15244001<br>10.5281/zenodo.15242188<br>10.5281/zenodo.15244003<br>10.5281/zenodo.15244005<br>10.5281/zenodo.15244007<br>10.5281/zenodo.15244009 |
| Fig. S1 | 10.5281/zenodo.17599004 |
| Fig. S2 | 10.5281/zenodo.17599254 |
| Fig. S3 | 10.5281/zenodo.17599590 |
| Fig. S4 | 10.5281/zenodo.17600219<br>10.5281/zenodo.17645734 |
| Fig. S5 | 10.5281/zenodo.15241738<br>10.5281/zenodo.15241583<br>10.5281/zenodo.15241628 |

**Data S1. Sequence of recombinant dCas9 proteins**

>dCas9-sfGFP-mNeonGreen

MGSSKRPAATKKAGQAKKKKGGSKRPAATKKAGQAKKKKEFGIHGVPAADKKYSIGLAIG
TNSVGWAVITDEYKVPSKKFKVLGNTDRHSIKKNLIGALLFDSGETAEATRLKRTARRRY
TRRKNRICYLQEIFSNEMAKVDDSFFHRLEESFLVEEDKKHERHPIFGNIVDEVAYHEKY
PTIYHLRKKLVDSTDKADLRLIYLALAHMIKFRGHFLIEGDLNPDNSDVDKLFIQLVQTY
NQLFEENPINASGVDAKAILSARLSKSRRLENLIAQLPGEKKNGLFGNLIALSLGLTPNF
KSNFDLAEDAKLQLSKDTYDDDLDNLLAQIGDQYADLFLAAKNLSDAILLSDILRVNTEI
TKAPLSASMIKRYDEHHQDLTLLKALVRQQLPEKYKEIFFDQSKNGYAGYIDGGASQEEF
YKFIKPILEKMDGTEELLVKLNREDLLRKQRTFDNGSIPHQIHLGELHAILRRQEDFYPF
LKDNREKIEKILTFRIPYYVGPLARGNSRFAWMTRKSEETITPWNFEEVVDKGASAQSFI
ERMTNFDKNLPNEKVLPKHSLLYEYFTVYNELTKVKYVTEGMRKPAFLSGEQKKAIVDLL
FKTNRKVTVKQLKEDYFKKIECFDSVEISGVEDRFNASLGTYHDLLKIIKDKDFLDNEEN
EDILEDIVLTLTLFEDREMIEERLKTYAHLFDDKVMKQLKRRRYTGWGRLSRKLINGIRD
KQSGKTILDFLKSDGFANRNFMQLIHDDSLTFKEDIQKAQVSGQGDSLHEHIANLAGSPA
IKKGILQTVKVVDELVKVMGRHKPENIVIEMARENQTTQKGQKNSRERMKRIEEGIKELG
SQILKEHPVENTQLQNEKLYLYYLQNGRDMYVDQELDINRLSDYDVDAIVPQSFLKDDSI
DNKVLTRSDKNRGKSDNVPSEEVVKKMKNYWRQLLNAKLITQRKFDNLTKAERGGLSELD
KAGFIKRQLVETRQITKHVAQILDSRMNTKYDENDKLIREVKVITLKSKLVSDFRKDFQF
YKVREINNYHHAHDAYLNAVVGTALIKKYPKLESEFVYGDYKVYDVRKMIAKSEQEIGKA
TAKYFFYSNIMNFFKTEITLANGEIRKRPLIETNGETGEIVWDKGRDFATVRKVLSMPQV
NIVKKTEVQTGGFSKESILPKRNSDKLIARKKDWDPKKYGGFDSPTVAYSVLVVAKVEKG
KSKKLKSVKELLGITIMERSSFEKNPIDFLEAKGYKEVKKDLIIKLPKYSLFELENGRKR
MLASAGELQKGNELALPSKYVNFLYLASHYEKLKGSPEDNEQKQLFVEQHKHYLDEIIEQ
ISEFSKRVILADANLDKVLSAYNKHRDKPIREQAENIIHLFTLTNLGAPAAFKYFDTTID
RKRYTSTKEVLDATLIHQSITGLYETRIDLSQLGGDGSPKKKRKVEDPKKKRKVDVRKGE
ELFTGVVPILVELDGDVNGHKFSVRGEGEGDATNGKLTLKFICTTGKLPVPWPTLVTTLT
YGVQCFARYPDHMKQHDFFKSAMPEGYVQERTISFKDDGTYKTRAEVKFEGDTLVNRIEL
KGIDFKEDGNILGHKLEYNFNSHNVYITADKQKNGIKANFKIRHNVEDGSVQLADHYQQN
TPIGDGPVLLPDNHYLSTQSVLSKDPNEKRDHMVLLEFVTAAGITHGMDELYKGSASGGG
GTSGGGSGSMVSKGEEDNMASLPATHELHIFGSINGVDFDMVGQGTGNPNDGYEELNLKS

TKGDLQFSPWILVPHIGYGFHQYLPYPDGMSPFQAAMVDGSGYQVHRTMQFEDGASLTVNYRYTYEGSHIKGEAQVKGTGFPADGPVMTNSLTAADWCRSKKTYPNDKTIISTFKWSYTTGNGKRYRSTARTTYTFAKPMAANYLKNQPMYVFRKTELKHSKTELNFKEWQKAFTDVMGMDELYKASGGGTYKGSSPAAKRVKLDGGSPAAKRVKLDLEHHHHHH

>dCas9-2xmScarlet

MGSSKRPAATKKAGQAKKKKGGSKRPAATKKAGQAKKKKEFGIHGVPAADKKYSIGLAIGTNSVGWAVITDEYKVPSKKFKVLGNTDRHSIKKNLIGALLFDSGETAEATRLKRTARRRYTRRKNRICYLQEIFSNEMAKVDDSFFHRLEESFLVEEDKKHERHPIFGNIVDEVAYHEKYPTIYHLRKKLVDSTDKADLRLIYLALAHMIKFRGHFLIEGDLNPDNSDVDKLFIQLVQTYNQLFEENPINASGVDAKAILSARLSKSRRLENLIAQLPGEKKNGLFGNLIALSLGLTPNFKSNFDLAEDAKLQLSKDTYDDDLDNLLAQIGDQYADLFLAAKNLSDAILLSDILRVNTEITKAPLSASMIKRYDEHHQDLTLLKALVRQQLPEKYKEIFFDQSKNGYAGYIDGGASQEEFYKFIKPILEKMDGTEELLVKLNREDLLRKQRTFDNGSIPHQIHLGELHAILRRQEDFYPFLKDNREKIEKILTFRIPYYVGPLARGNSRFAWMTRKSEETITPWNFEEVVDKGASAQSFIERMTNFDKNLPNEKVLPKHSLLYEYFTVYNELTKVKYVTEGMRKPAFLSGEQKKAIVDLLFKTNRKVTVKQLKEDYFKKIECFDSVEISGVEDRFNASLGTYHDLLKIIKDKDFLDNEENEDILEDIVLTLTLFEDREMIEERLKTYAHLFDDKVMKQLKRRRYTGWGRLSRKLINGIRDKQSGKTILDFLKSDGFANRNFMQLIHDDSLTFKEDIQKAQVSGQGDSLHEHIANLAGSPAIKKGILQTVKVVDELVKVMGRHKPENIVIEMARENQTTQKGQKNSRERMKRIEEGIKELGSQILKEHPVENTQLQNEKLYLYYLQNGRDMYVDQELDINRLSDYDVDAIVPQSFLKDDSIDNKVLTRSDKNRGKSDNVPSEEVVKKMKNYWRQLLNAKLITQRKFDNLTKAERGGLSELDKAGFIKRQLVETRQITKHVAQILDSRMNTKYDENDKLIREVKVITLKSKLVSDFRKDFQFYKVREINNYHHAHDAYLNAVVGTALIKKYPKLESEFVYGDYKVYDVRKMIAKSEQEIGKATAKYFFYSNIMNFFKTEITLANGEIRKRPLIETNGETGEIVWDKGRDFATVRKVLSMPQVNIVKKTEVQTGGFSKESILPKRNSDKLIARKKDWDPKKYGGFDSPTVAYSVLVVAKVEKGKSKKLKSVKELLGITIMERSSFEKNPIDFLEAKGYKEVKKDLIIKLPKYSLFELENGRKRMLASAGELQKGNELALPSKYVNFLYLASHYEKLKGSPEDNEQKQLFVEQHKHYLDEIIEQISEFSKRVILADANLDKVLSAYNKHRDKPIREQAENIIHLFTLTNLGAPAAFKYFDTTIDRKRYTSTKEVLDATLIHQSITGLYETRIDLSQLGGDGSPKKKRKVEDPKKKRKVDMVSKGEAVIKEFMRFKVHMEGSMNGHEFEIEGEGEGRPYEGTQTAKLKVTKGGPLPFSWDILSPQ

FMYGSRAFTKHPADIPDYYKQSFPEGFKWERVMNFEDGGAVTVTQDTSLEDGTLIYKVKL
RGTNFPPDGPVMQKKTMGWEASTERLYPEDGVLKGDIKMALRLKDGGRYLADFKTTYKAK
KPVQMPGAYNVDRKLDITSHNEDYTVVEQYERSEGRHSTGGMDELYKGSASGGGGTSGGG
SGSMVSKGEAVIKEFMRFKVHMEGSMNGHEFEIEGEGEGRPYEGTQTAKLKVTKGGPLPF
SWDILSPQFMYGSRAFTKHPADIPDYYKQSFPEGFKWERVMNFEDGGAVTVTQDTSLEDG
TLIYKVKLRGTNFPPDGPVMQKKTMGWEASTERLYPEDGVLKGDIKMALRLKDGGRYLAD
FKTTYKAKKPVQMPGAYNVDRKLDITSHNEDYTVVEQYERSEGRHSTGGMDELYKASGGG
TYKGSSPAAKRVKLDGGSPAAKRVKLDLEHHHHHH
